# Chewing Through Challenges: Exploring the Evolutionary Pathways to Wood-Feeding in Insects

**DOI:** 10.1101/2023.12.27.573094

**Authors:** Cristian F. Beza-Beza, Brian M. Wiegmann, Jessica A. Ware, Matt Petersen, Nicole Gunter, Marissa E. Cole, Melbert Schwarz, Matthew A. Bertone, Daniel Young, Aram Mikaelyan

**Affiliations:** Department of Entomology, University of Minnesota, St Paul, MN, USA; Department of Entomology and Plant Pathology, North Carolina State University, Raleigh, NC, United States; Department of Biological Sciences, Rutgers University, Newark, NJ 08854, USA; Biodiversity and Geosciences Program, Queensland Museum, South Brisbane, Queensland, Australia; Department of Entomology, University of Wisconsin, Madison, WI 53706

**Author notes:** Correspondence: Cristian F. Beza-Beza, Aram Mikaelyan.

## Abstract

Decaying wood, while an abundant and stable resource, presents considerable nutritional challenges due to its structural rigidity, chemical recalcitrance, and low nitrogen content. Despite these challenges, certain insect lineages have successfully evolved saproxylophagy (consuming and deriving sustenance from decaying wood), impacting nutrient recycling in ecosystems and carbon sequestration dynamics. This study explores the uneven phylogenetic distribution of saproxylophagy across insects and delves into the evolutionary origins of this trait in disparate insect orders. Employing a comprehensive analysis of gut microbiome data, encompassing both previously published datasets and newly generated data, from both saproxylophagous insects and their non-saproxylophagous relatives, this *Hypothesis* paper discusses the broader phylogenetic context and potential morphological, physiological, and symbiotic adaptations necessary for this dietary specialization. The study proposes the “Detritivore-First Hypothesis,” suggesting an evolutionary pathway to saproxylophagy through detritivory, and highlights the critical role of symbiotic gut microbiomes in the digestion of decaying wood. The article aims to provide a deeper understanding of the macroevolutionary landscape and mechanisms underpinning the multiple origins and distribution of saproxylophagy in insects.

## Evolutionary Pathways to Saproxylophagy

Saproxylophagy, or the capacity to feed on decaying wood, offers an intriguing case study in the evolution of intricate adaptations to highly demanding diets. While decaying wood is both an abundant and a relatively stable resource across ecosystems, it poses significant nutritional challenges to those organisms that depend on it ^[1]^. Wood consists predominantly of cellulose, hemicellulose, and lignin organized in distinct secondary structures with cellulose fibrils embedded in a hemicellulose matrix intertwined with lignin ^[2]^. The resulting structural rigidity, chemical recalcitrance, and low nitrogen content make it virtually impossible for most animal species to subsist on wood. Yet, in the face of these formidable odds, a few major lineages of insects have successfully evolved saproxylophagy ^[2]^. This ability has enormous consequences for both the recycling of nutrients in diverse ecosystems as well as for the dynamics of carbon sequestration, among other geochemical and ecological phenomena that have had lasting implications for our civilization and the planet. Indeed, coal was formed when carboniferous trees died and failed to decompose. This occurred during a time early in the evolution of lignin- and cellulose-degrading organisms, at least 100 million years before the evolution of wood-degrading fungi^[3]^, passalid beetles^[4]^ and termites^[5]^.

While saproxyly (in association in decaying wood) is relatively common among insects ^[6]^, saproxylophagy has evolved in only four orders of insects out of 30, notably Coleoptera (beetles ^[7–9]^), Blattodea (cockroaches and termites ^[10,11]^), Diptera (flies^[12]^) and Lepidoptera (moths ^[13]^). Clearly, lineages with different phylogenetic contexts have the capacity to evolve this dietary specialization. However, although this feeding habit has emerged in evolutionarily very distinct insect orders, its phylogenetic distribution is far from uniform. First, the lineages that have evolved saproxylophagy are phylogenetically distant from each other; Blattodea are non-holometabolous members of the Polyneoptera while the remaining three saproxylophagous lineages are dispersed across the Holometabola tree of life. Second, even within specific orders, saproxylophagous lineages are not randomly distributed. Thus, saproxylophagy raises a suite of compelling evolutionary questions about its origins. If the digestion of decaying wood is a complex trait, how has it evolved across unrelated insect lineages? If only a few insect lineages have been capable of evolving saproxylophagy, what kinds of dietary specializations might predispose certain clades to develop this challenging feeding habit?

The phylogenetic distribution of saproxylophagy across insects suggests that the evolutionary contexts, costs and benefits of this diet may vary significantly across clades. Furthermore, this apparent unequal scattering of saproxylophagy points to the existence of specific ecological niches and evolutionary forces that might enable certain insect lineages to adopt wood-feeding habits more easily than others might. For example, lineages that, due to their ecological, geographical, or historical circumstances, find themselves in “favored niches” rich in decaying wood, can adapt to exploit an abundant, albeit nutrient-poor, resource. On the other hand, ecological opportunity alone might not be sufficient for the evolution of saproxylophagy; clades can differ in the extent to which they possess the “favored” genotypic and phenotypic attributes necessary for wood-feeding. Within this framework, it is imperative to consider the symbiotic nature of decaying wood digestion in insects, and regard the microbial symbionts associated with digestion as vital extensions of these favored physiological attributes^[1]^.

We hypothesize that the evolution of saproxylophagy entailed a cascade of correlated changes spanning anatomical, physiological, and microbial dimensions, each pivotal in addressing the nutritional challenges posed by decaying wood. These evolutionary trajectories likely reflect adaptations to both ecological opportunities presented by favored niches, as well as biological capacities intrinsic to specific lineages in the form of favored attributes. Employing a macroevolutionary approach to understand saproxylophagy in insects may elucidate the forces underpinning this challenging dietary specialization. In this Hypothesis piece, we discuss the broader ecological and phylogenetic context surrounding saproxylophagous clades and its implications for their origins. We consider the feeding habits of relatives of saproxylophagous clades, and the potential morphological and physiological adaptations necessary to overcome the challenges associated with relying on a recalcitrant diet. Our dissection of saproxylophagy recognizes the microbiome as a key mediator, integrating insect physiology and evolutionary strategies with the ecological opportunities that drive adaptation. Our arguments are backed by comprehensive analyses of both published and newly generated gut microbiome data from saproxylophagous insects and their non-saproxylophagous relatives. Furthermore, to highlight the tight symbiotic relationship between decaying wood feeders and their associated microbes, we also present a phylogenetic analysis of a widely distributed lignocellulose-degrading bacterial lineage.

We conclude by elaborating the “Detritivore-First Hypothesis”, suggesting a potentially generalizable evolutionary pathway to saproxylophagy through detritivory. This hypothesis is informed by the following observations: (1) the close phylogenetic proximity of detritivorous and saproxylophagous lineages, (2) the presence of morphological and physiological traits in detritivore insects that could be considered preadaptations facilitating the evolution of saproxylophagy, (3) the reliance on, and the similar composition of, complex microbiomes in detritivorous and saproxylophagous insects, and (4) the shared distribution and close phylogenetic relationship of fiber-digesting microbes found in both detritivorous and saproxylophagous insects.

## Macroevolutionary Landscape and Mechanisms Underpinning the Multiple Origins and Unequal Distribution of Saproxylophagy

Several explanations for the relatively rare and scattered distribution of saproxylophagy across Insecta may exist, but current phylogenetic and ecological considerations narrow the list of plausible options:

Although all extant insect orders have been exposed to decaying wood, only a few lineages have been exposed to the adaptive landscape, in the contexts of the details of their niches, that would have favored the evolution of a diet based on decaying wood. For example, within herbivore lineages, those that have interacted and consumed cambium may be prone to evolving saproxylophagy, in particular. Adaptations to utilize lignocellulose-rich material combined with selective pressures like those posed by the challenges of the plant’s defense mechanisms could facilitate the transition to feeding on decaying wood. Although feeding on cambium and other living tissues is challenging due to a complex array of plant defense strategies, wood of living plants is less recalcitrant and relatively more nutritious than decaying wood ^[14]^.

Another niche that might incline species towards saproxylophagy is feeding on fungus that grows on it (xylomycophagy); although the two feeding habits provide overlapping favored niches, the fundamental differences in the composition of fungi and wood each demand distinct physiological and enzymatic capabilities, making this particular transition seem unlikely. Gimmel and Ferro (2018) noted that beetles dwelling and feeding on “wood-rotting fungal bodies” constitute “the largest single category of saproxylic beetles”, and even when some of the families associated with this feeding habit are species rich, there are no known transitions to wood-feeding. This suggests that obligate fungus feeders have unique traits that do not inherently facilitate a shift to true wood feeding.

Alternatively, insects specialized in the decomposition of rotting plant material (e.g., leaf litter) composed of limited and variable amounts of decaying wood provide the right adaptive landscape to specialize in the consumption of substrates with a gradual increase in amounts of decaying wood. With either of these scenarios, protosaproxylophagous insects likely possessed specific traits that facilitate the evolution of saproxylophagy, i.e. preadaptations. These include anatomical (e.g., robust mandibles for grinding, the potential for hindgut enlargement and compartmentalization), physiological (e.g., intestinal pH, and the recruitment of complex microbiomes capable of fiber digestion and fermentation), or behavioral (e.g., trophallaxis to provide young with predigested wood, or to reinoculate guts with a full complement of the microbiome) traits that provide a foundation for the evolution of saproxylophagy.

The taxonomic boundaries of saproxylophagy are best understood for the cockroach (Blattodea) lineage Kittrickea *sensu* Evangelista et al. 2019 ^[5]^ and the beetle (Coleoptera) family Passalidae. The primordially saproxylophagous Kittrickea comprises the subsocial woodroaches of the family Cryptocercidae and the eusocial termites ^[5]^. The beetles of the family Passalidae (bess beetles) are specialized wood-feeders and likely are the sole representatives of Coleoptera, where the adults also thrive on wood. In contrast, for other saproxylophagous insect lineages, like the Blaberidae (Blattodea), Scarabaeidae (Coleoptera), Bibionomorpha (Diptera), Tipulidae (Diptera), and Limoniidae (Diptera), the taxonomic boundaries for saproxylophagy remain uncertain.

Current phylogenetic knowledge reveals two distinct patterns of saproxylophagous insects. The first is that of saproxylophagous species are found in large monophyletic clusters, suggesting a shared ancestral origin of wood-feeding and a pronounced tendency for niche conservatism (e.g., Kittrickea, Panesthiinae, Passalidae, Ctenophorinae) (Figs 1a-b). In contrast to these deeper origins of saproxylophagy, lineages like Blaberidae, Scarabaeidae and Limoniidae likely contain multiple independent origins, indicating a more flexible or dynamic tendency for trophic specialization and/or its loss through extinction of saproxylophagous lineages (Figs 1c-e).

**Fig. 1.**
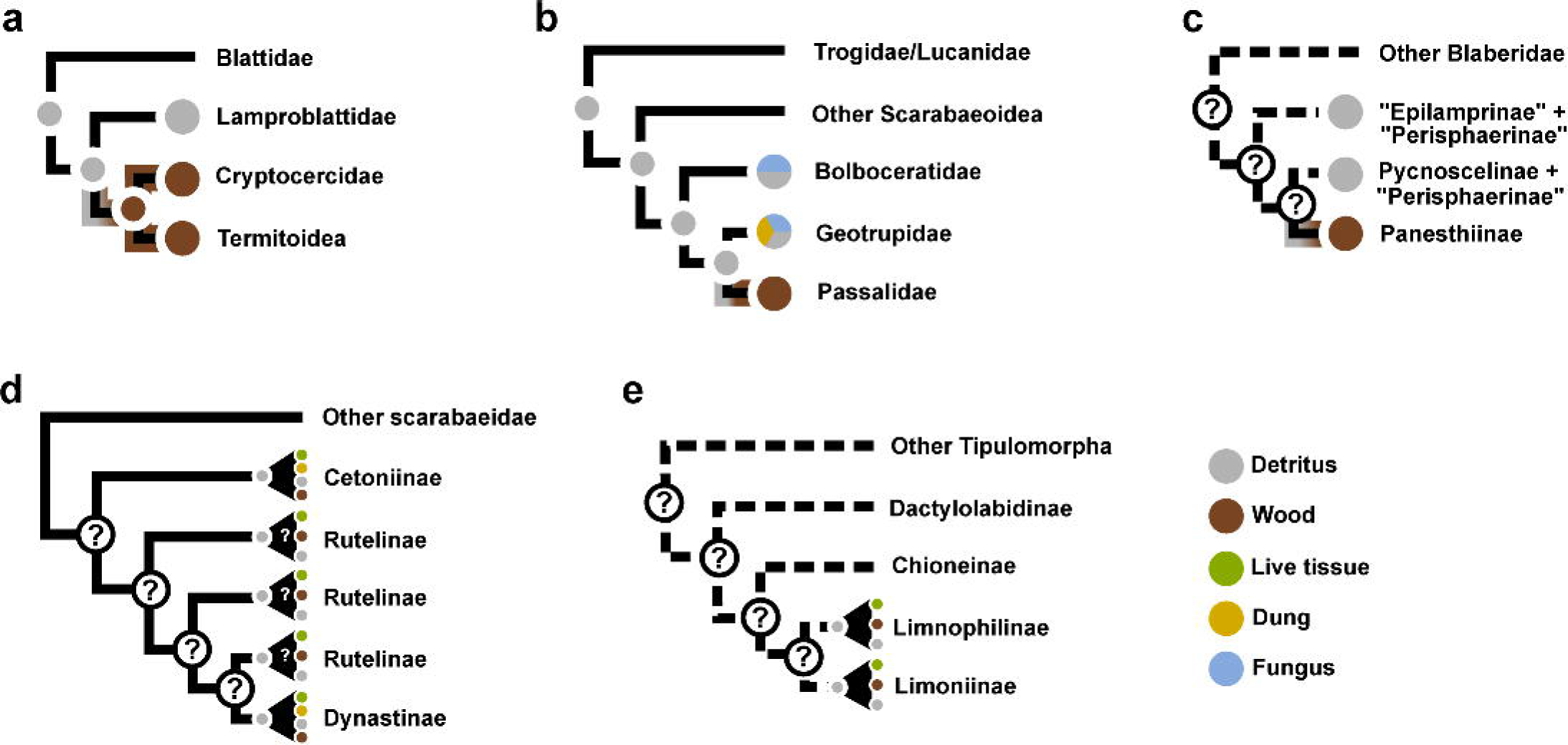
Phylogenetic Overview of decaying wood Feeding Lineages and Potential Transition to Saproxylophagy. This Fig. illustrates the possible evolutionary transitions to saproxylophagy in various insect lineages; **a)** Cryptocercidae and Termitoidea, **b)** Passalidae, **c)** Blaberidae, **d)** Scarabaeidae, and **e)** Limoniidae. Circles on the tips depict the feeding habits of extant lineages, while circles on the nodes represent hypothetical ancestral feeding habits

In some cases (e.g. Kittrickea, Blaberidae, and Passalidae), it is clear that the transition to saproxylophagy was derived from detritivory (Figs 1a-c). The most comprehensive study on Blattodean phylogeny (Evangelista et al. 2019) establishes Kittrickea as sister to Lamproblattidae (Fig. 1a). For Blaberidae, although the taxonomic boundaries of saproxylophagy are unknown, the decaying wood feeding Panesthiinae subfamily and the Zetoborinae genus *Parasphaeria* are nested within this predominantly detritivorous clade (Fig. 1c)^[15,16]^. Passalidae, which represents one of the oldest origins of saproxylophagy ^[4,17]^ is sister to Geotrupidae, which is predominantly detritivorous ^[18,19]^ (Fig. 1c). The previous examples highlight the evident phylogenetic relationship between saproxylophagous and detritivorous insects. In other saproxylophagous lineages (e.g, Scarabaeidae and Limoniidae), detritivory is common and may be considered the ancestral condition, however this is yet unconfirmed through detailed phylogenetic study. Ancestral detritivory in these groups supports the idea that these detritivorous clades possessed both the ecological opportunity and potential pre-adaptations necessary for saproxylophagy.

Detailed phylogenetic studies will be key to unraveling the trophic history for diverse saproxylophagous lineages. For example, in Scarabaeidae (Coleoptera) and Limoniidae (Diptera) which are large, mainly detritivorous families, larvae are found in, near, and around decaying wood. These lineages also have a large number of species associated with other feeding habits, including fungivory, detritivory, carnivory, omnivory, and phytophagy. The subfamilies Rutellinae, Dynastinae, and Cetoniinae within Scarabaeidae demonstrate recent and multiple origins of larval saproxylophagy, contrasting with their relatives that feed on varied diets, including dung, carrion, fungi, leaf litter, soil, and roots ^[20]^ (Fig. 1d). Similarly, among the flies, the association of numerous species within Bibionomorpha, Chironomidae, Limoniidae, and Tipulidae has led to the speculation that they are indeed wood consumers ^[21]^. However, the most compelling evidence of obligate wood-feeding comes from the limoniid crane flies, with at least 25 genera recorded from dead wood ^[22]^, including *Limonia* ^[23]^ and *Lipsothrix* ^[24,25]^ in which consumption of wood has been directly demonstrated. Although the phylogeny is still poorly resolved ^[22]^, the broad occurrence of saproxylophagy observed in these taxa alone suggest multiple independent phylogenetic origins arising from lineages primarily feeding on decomposing plant material (Fig. 1e).

Macroevolutionary studies addressing the estimation of ancestral feeding habits within groups like Scarabaeidae and Limoniidae will provide valuable insights for hypotheses testing on possible paths for the evolution of saproxylophagy, beyond just detritivory. Complementary to better phylogenies, highly resolved quantitative analyses of diets for saproxylophagous insects and their non-saproxylophagous relatives are necessary. Currently, even the most advanced gut content analyses, like the methods developed by Donovan et al. ^[26]^ for categorizing termites into specific lignocellulose-feeding groups, are prone to subjective interpretation, due to factors such as observer-dependent visual analysis, limitations of microscopy resolution, and challenges in replicability. This situation underscores the need for more refined methods that can clearly ascribe the diet of any insect species to the humification spectrum of lignocellulose. These data can reveal a better understanding of the factors that drive not only the evolution of saproxylophagy, but also the mechanisms of trophic specialization within these diverse clades.

## Anatomical and Physiological Adaptations to Wood-Feeding

Investigating adaptations that allowed the evolution of saproxylophagy may reveal key evolutionary starting points and common pathways to wood-feeding in these lineages. However, the traits in question cannot simply be considered in isolation. Building on Kemp’s concept of “correlated progression”^[27]^, traits associated with feeding, processing food, nutrition, and excretion are intricately interdependent. A change in one attribute, like mouthpart adaptations, necessitates concurrent alterations in other attributes, ensuring evolutionary coherence, and promoting the evolution of integrated innovations. For instance, the evolution of robust mandibles for cutting and grinding wood would need to be accompanied by changes, such as the elongation or dilation of specific regions of the gut to accommodate and process the increased intake of fibrous material.

Digestive efficiency of animals is also inextricably linked to both gut retention time and the distribution of digestive enzymes ^[28]^. Consequently, morphological adaptations contributing to gut compartmentalization, along with physiological adaptations such as alterations in digestive enzyme activity, can serve as valuable indicators of feeding ecology. Adaptations exhibited by mammals to high-fiber diets can be especially instructive in this context, where we observe two significant traits: (1) expansion of the luminal volume and compartmentalization in either the foregut (as seen in the ruminants’ rumen) or the hindgut (as exemplified by the presence of haustra, taeniae, and mucosal folds in monogastric ungulates); and (2) a marked reduction in salivary gland tissue, reflecting a heightened reliance on the microbiome for enzymatic contributions ^[29]^. In both cases, these evolutionary adaptations serve to maximize the residence time of recalcitrant plant material within specific gut compartments, thereby facilitating microbial fermentation and increasing surface area for the absorption of fermentation products.

Saproxylophagous insects appear to have undergone a process analogous to the evolution of herbivory in mammals. These insects have developed a unique set of morphological adaptations to cope with the challenges of a recalcitrant lignocellulose-based diet. All described wood-feeding insect lineages among the tutricablattids (Isoptera and Cryptocercidae) ^[30–32]^, panesthiine cockroaches ^[33–35]^, and passalid beetles ^[36–38]^ rely heavily on hindgut fermentation for lignocellulose digestion. For this reason, we expect that wood-feeding insects should possess significantly larger and more compartmentalized hindguts, compared to non-wood-feeding insects. This increase in relative hindgut volume and compartmentalization appears to correspond with the maintenance of distinct physicochemical microenvironments (such as pH, redox potential, and H_2_/O_2_ partial pressure) along the gut axis ^[30,33,35,37]^.

While the above-mentioned morphological adaptations provide insights into the symbiotic digestion of wood, enzymatic activity across various gut regions further refines our understanding of these processes. Indeed studies on termites have provided invaluable insights into the distribution of enzyme activity across gut sections ^[1]^. However, a mere quantification of enzyme activity across different compartments is likely to overlook the contributions of varying residence times of food particles in different compartments. For instance, in the wood-feeding blaberids *Salganea taiwanensis* and *Panesthia angustipennis*, almost half of the cellulose and xylan activity is associated with the crop, whereas only about a fifth is linked to the hindgut ^[34]^. However, considering the vast difference in residence times observed even in detritivorous *Periplaneta americana*, where particles remain in the hindgut for over 18 hours compared to less than 30 minutes in the crop ^[39]^, it becomes essential to take a more nuanced look at how enzyme activity is not just distributed across different gut regions, but also optimized in the context of residence times. Extended residence times within the gut not only enable the necessary enzymatic breakdown but also allow for the optimal extraction of nutrients from challenging substrates. For example, gut retention times likely play a pivotal role in the effective digestion of lignocellulose in wood-feeding roaches ^[34]^. Unlike the detritus-based diets predominant in many cockroaches, decaying wood is inherently more recalcitrant and nutrient-poor ^[40]^. Understanding the interplay between gut retention times and enzymatic activity is essential for deciphering the evolutionary adaptations that allow insects to thrive on wood-based diets.

Some of the above-mentioned morphological and physiological adaptations play a pivotal role in the maintenance of complex microbiomes ^[30,33,36,41]^. For instance, the gut regions, with their specific physicochemical conditions and residence times, create specialized environments that foster the presence and function of these microbial communities in termites, passalid beetles, and panesthiine cockroaches. Notably, germ-free cockroaches, devoid of their microbiomes, exhibit non-distended guts ^[42]^, highlighting the microbiome’s influence on gut morphology ^[43]^. Consequently, the insect gut is a coordinated ecosystem, where both the host and its symbionts work in tandem to efficiently break down wood and extract nutrients from lignocellulose. Importantly, it is the quintessentially holobiotic physiology of saproxylophagous insects, and the anatomical and physiological changes in the animal’s gut itself that enable the enrichment of fiber-fermenting microbes; it this fully integrated system that is inherited from detritivorous ancestors to enable fully specialized wood-feeding.

## Insects and Microbes: The Interlocking Gears of Symbiotic Wood Digestion

To better understand how saproxylophagy evolved in diverse lineages, it is valuable to evaluate how symbiotic relationships are established between insects and their associated microbes. Although saproxylophagous marine invertebrates have evolved endogenous enzymes employed in degrading lignocellulose ^[44]^, the mechanism to digest decaying wood in insects resembles a sequential bioreactor ^[45]^. It heavily relies on the gut microbiome, meshing the insect’s own adaptations with the contributions of microbial symbionts to achieve an efficient extraction of nutrients from this complex resource. In the context of the differences among these “bioreactors”, and their morphological “infrastructure”, we can answer specific questions about the formation of such complex symbioses in the evolution of insects, such as, could the fiber-digesting symbionts of saproxylophagous insects have been environmentally acquired or were they inherited from their protosaproxylophagous ancestors?

Since most metazoans lack the inherent capacity to digest lignocellulose independently, insect saproxylophagy presents a compelling system to study the role of microbial symbioses in facilitating the emergence of key innovations, expanding ecological niches, and driving diversification. However, these symbioses stand on coordinated platforms of physiological, anatomical, and behavioral adaptations in the host ^[1]^. In order to be metabolically feasible, the establishment of digestive symbioses was a necessary step towards unlocking the digestion of decaying wood. Insects digest wood through a complex interplay between the insect and its associated microbiome. This process typically begins with the physical breakdown of wood by the insect’s sclerotized mandibles and partial biochemical degradation by the insect’s endogenous enzymes ^[1]^. The wood fibers then flow into an enlarged portion of the hindgut that acts as the primary fermentation chamber, where they are degraded by a specific microbial community ^[31,38]^.

The hindgut of insects that consume plant material, such as wood, litter, and forms of humified lignocellulose possesses adaptations that maximize microbial degradation and fermentation of food particles to short-chain fatty acids ^[46]^. Wood-feeding insects, including termites, cockroaches and passalid beetles, are all characterized by a distended hindgut, that serves as the primary fermentation chambers for the breakdown of wood particles ^[30,33,35,38,47]^. Despite their relatively small volumes, these hindguts are unexpectedly complex ecosystems, harboring some of the most diverse insect gut microbiomes ^[48]^. The factors contributing to this complexity are still poorly understood, but the strong connections between the microheterogeneity observed in these hindguts, microbiome composition, and lignocellulose digestion are evident in termites ^[30,31,41,49–51]^, cockroaches ^[33,35]^, and passalid beetles ^[36–38]^.

The surfaces of wood fibers in the insect hindgut are central to the process of lignocellulose digestion, serving as microhabitats for colonization by fiber-associated bacteria ^[31,34,38]^. Although every wood-feeding insect clade displays convergent evolution in their reliance on fiber-associated microbiomes, there is marked distinctiveness in their bacterial compositions. For instance, the fibers in the hindgut of the passalid beetle *Odontotaenius disjunctus* are consistently associated with Firmicutes ^[38]^, whereas those in the termites *Nasutitermes corniger* and *Nasutitermes takasagoensis*. are predominantly colonized by Fibrobacterota and Spirochaetota ^[31]^. These compositional differences underscore the specialized recruitment and co-evolution with distinct sets of fiber-digesting bacteria. However, a shift is observed in more recent research exploring saproxylophagous cockroaches ^[35,52]^ and passalid beetles ^[38]^, which reveals a more universal core microbiome, transcending taxonomic boundaries but still restricted to lignocellulose-feeding insects. Highlighting this, our 16S-rRNA-based phylogenetic analysis (Fig. 2; Supplementary Files 1 and 2) of the putatively fiber-associated Christensenellaceae R-7 group shows multiple insect-specific clades with representatives from lignocellulose-feeding termites, cockroaches, scarab beetles, and the wood-feeding limoniid crane fly *Epiphragma solatrix*.

**Fig. 2.**
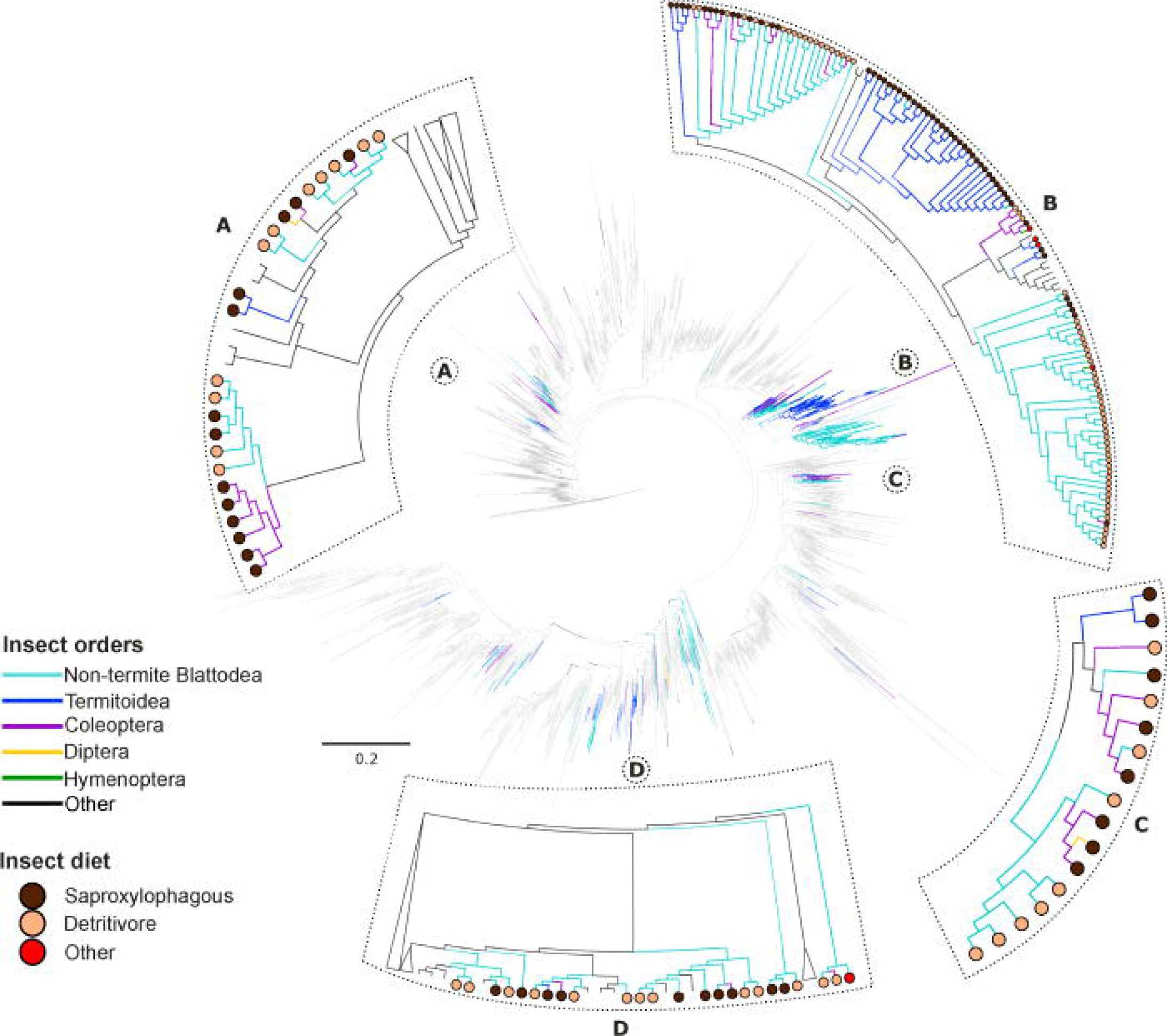
Phylogenetic Analysis of R-7 Group Bacterial Lineages within Christensenellaceae. This maximum likelihood tree showcases the bacterial lineages of the R-7 group. Clades A, B, and C predominantly indicate bacterial lineages associated with insect hosts. Branch colors depict the host association, while circles at the tips highlight the respective host’s diet. See Supplementary Information for the sequence alignment and phylogenetic tree file.

Given the critical role of complex gut microbiomes in wood digestion, it is useful to understand the insect along with its fiber-digesting symbionts as a single physiological and evolutionary unit, commonly referred to across animal systems as the holobiont ^[53]^. Highly similar distributions of these fiber-digesting bacteria are found among detritivorous relatives, as is observed in the relationship between Fibrobacterota lineages from wood-feeding termites and early-branching litter-feeding cockroaches ^[54]^. Our findings indicate that litter feeders and wood feeders exhibit a notably higher congruence in community structure (at the bacterial genus level) as compared to how wood feeders’ overlap with other dietary groups, such as those feeding on live wood (Fig. 3). While much of our understanding of this shared microbiome similarity has historically been anchored in studies of termites and cockroaches, our preliminary analysis extends this knowledge to diverse saproxylophagous groups. Specifically, patterns observed in the passalid beetle *O. disjunctus*, the telephone pole beetle *Micromalthus debilis*, and the limoniid *E. solatrix* reinforce this hypothesis. Such distribution patterns deserve more attention as they offer a window into the identity of the symbionts that were integral to lignocellulose digestion in protosaproxylophagous ancestors. Among the many adaptations that lead to saproxylophagy, the gut microbiome stands out as the most dynamic and plastic component, which is simultaneously also the preeminent contributor to the metabolic diversity needed to break down a resource as complex as lignocellulose. Therefore, the microbiome has the capability to most quickly adapt to variations in diet, simultaneously having the potential to shape the long-term evolutionary paths of these insects.

**Fig. 3.**
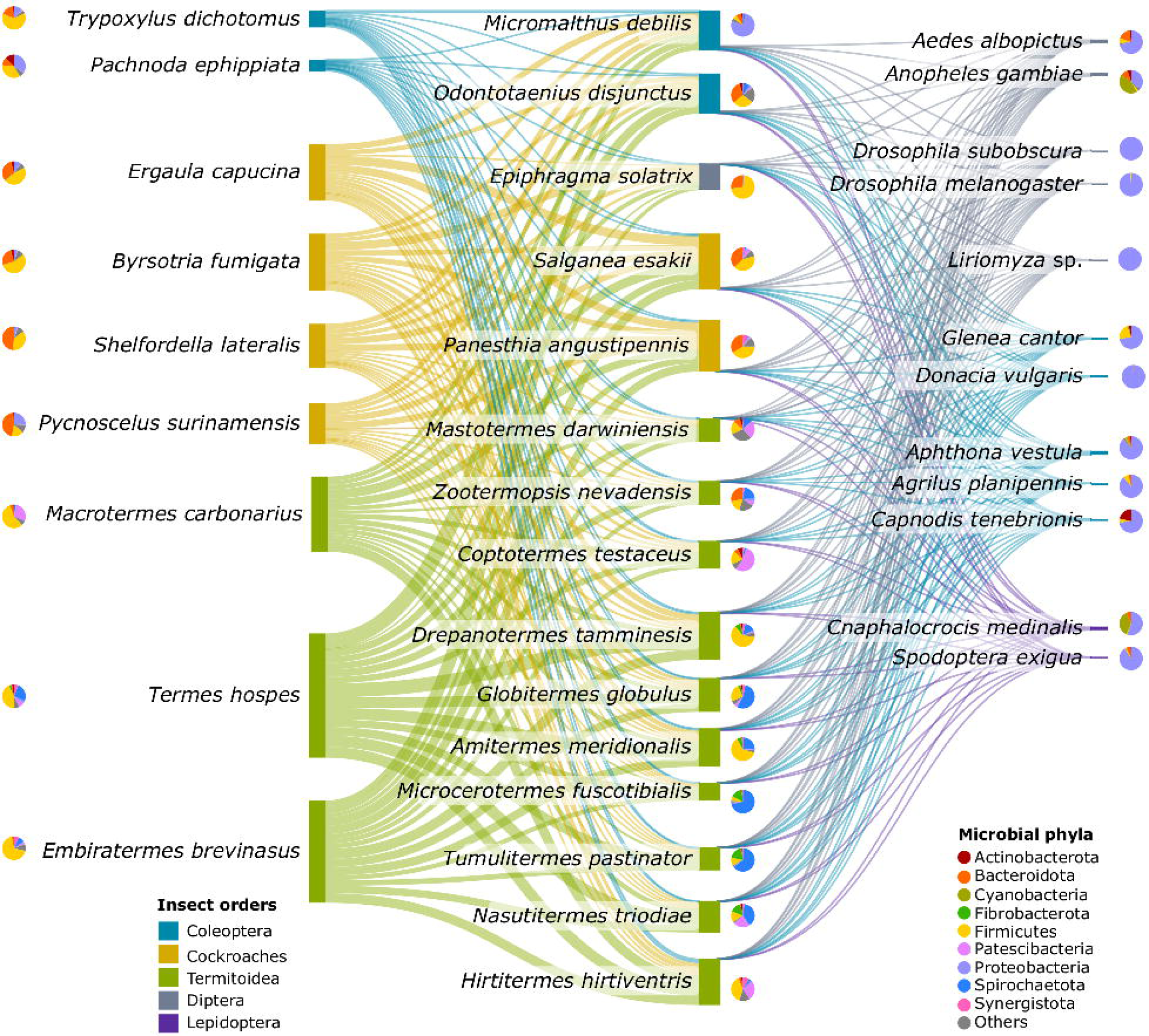
Microbiome Similarities Across Insect Diet Categories. This Sankey diagram visually represents the similarities in gut microbiome compositions across three primary insect diet categories. Column 1 features litter and humus feeders, Column 2 showcases wood feeders, and Column 3 highlights non-lignocellulose feeders. The diagram delineates comparisons between columns 1 and 2, as well as between columns 2 and 3. The width of the connecting flows is proportional to the Bray-Curtis similarities in their gut microbiomes.

## The Detritivore-First Hypothesis

In our synthesis of the observations discussed, we propose the “Detritivore-First” hypothesis. Detritivory encompasses the consumption of decaying organic materials, including those from both animal and plant origins ^[40]^. However, our focus is on specific insect groups that consume rotting plant material, which align with Groups I and II of the detritus-feeding saprophages as classified by Terra et al. ^[40]^ This includes insects that feed on dead plant matter and those that feed on feces but excludes those feeding on dead animal matter. Terra et al. ^[40]^ further indicate that, from a nutritional and chemical standpoint, animal dung is similar to decaying plant tissue, except that it is more humified and enriched with microbes. With the Detritivore-First hypothesis, we posit that detritivorous insects, through their unique dietary composition, possessed foundational preadaptations that facilitate the emergence of specialized symbiotic digestive systems that are now emblematic of all saproxylophagous insects (Fig. 4).

**Fig. 4.**
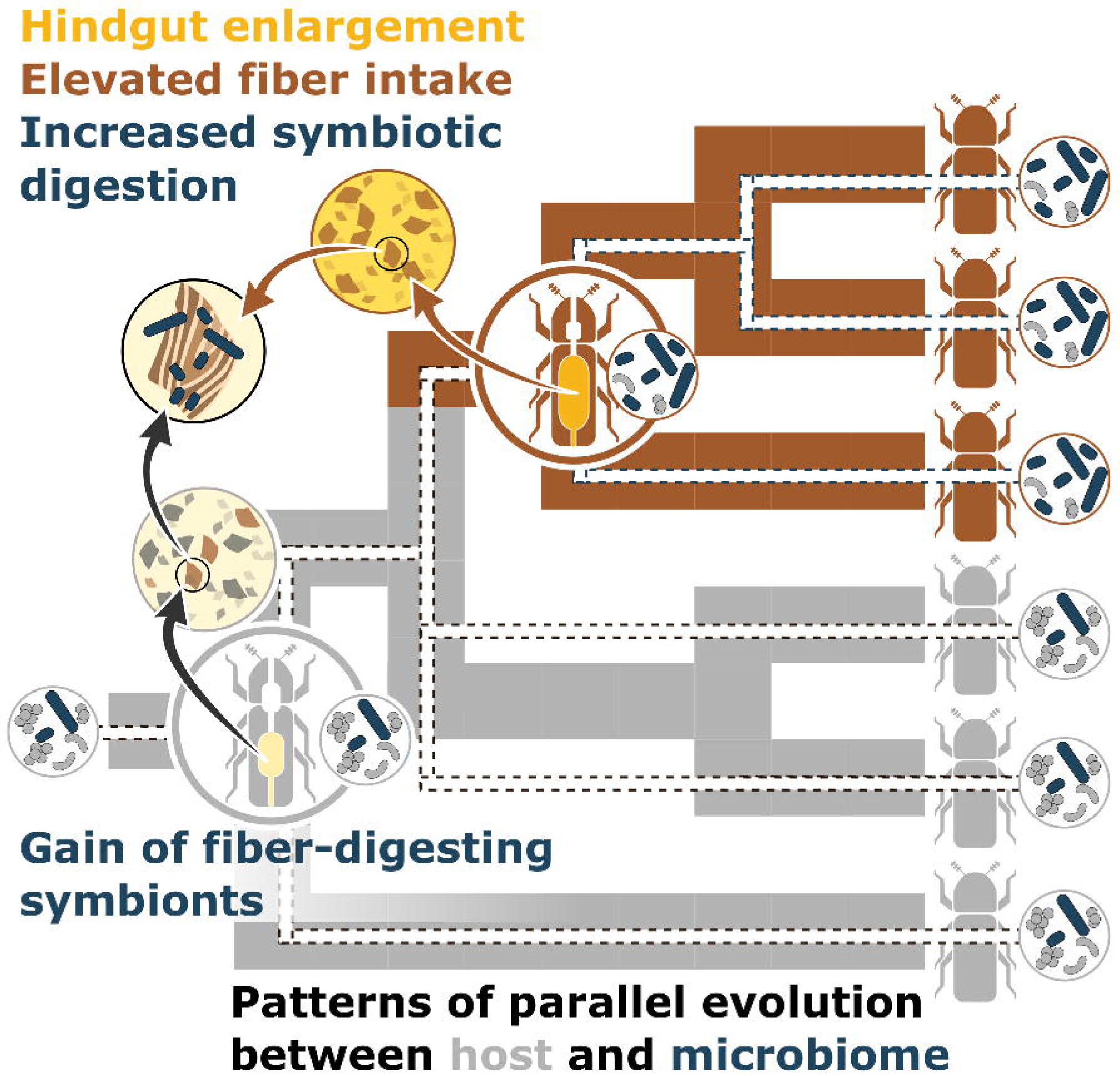
The Detritivore-First Hypothesis. A visualization of the hypothetical progression influenced by the coordinated changes in dietary fiber content, relative hindgut volume, and fiber-digesting symbionts. The interplay of these elements suggests a potential evolutionary pathway from detritivory to saproxylophagy.

Support for this notion is drawn from the following empirical evidence:

1. **Phylogenetic patterns:** Recognized wood-feeding insect clades are predominantly nested within broader detritivorous clades (Fig. 1). Notable examples include the Tutricablattae ^[5]^, and the Passalidae ^[17]^
2. **Morphophysiological parallels:** A striking similarity exists not only in the morphological and physiological traits between unrelated saproxylophagous clades (e.g. wood-feeding termites and passalid beetles), but also between those of their respective detritivorous relatives (e.g. detritivorous cockroaches and scarab beetles).
3. **Holobiotic dependency:** Both saproxylophagous and detritivorous insects are universally reliant on hindgut microbiomes. Interestingly, these microbiomes exhibit comparable compositions across species, as documented in studies on Blattodea ^[52,54]^ and Passalidae ^[38]^. (Fig. 3).
4. **Shared Symbiotic Relationships:** The shared distribution of fiber-digesting symbionts between saproxylophagous insects and their detritivorous counterparts suggests these microbes originated in the protosaproxylophagous ancestor. As wood fibers became more prevalent in the diets of saproxylophagous lineages, the abundance of these fiber-associated microbial lineages likely correspondingly increased, emphasizing the diet-driven changes in the gut microbiome of wood feeders. (Fig. 2)

If saproxylophagy is the result of the specialization along a continuum from decaying plant saprophagy in which detritivorous insects operate, and if the generalized model summarized in Fig. 4 holds true, then we can make the following predictions:

1. **Ecomorphological proximity:** Saproxylophagous lineages ought to be both phylogenetically and ecomorphologically close to detritivores consuming high-fiber diets.
2. **Compartmentalized guts:** Detritivorous relatives of saproxylophagous lineages should exhibit compartmentalized guts, including a hindgut fermentation chamber for the slow digestion of plant material.
3. **Hindgut expansion:** Although most detritivorous insects already have a distended hindgut, the transition from detritivory to saproxylophagy should be accompanied by a further expansion in its volume (relative to the total volume of the gut). This expansion likely leads to extended retention times, optimizing microbial digestion and fermentation.
4. **Microbial Phylogenetic continuity:** Detritivorous relatives ought to harbor a fiber-associated microbiome consisting of bacterial lineages closely related to those present in saproxylophagous insects. This suggests evolutionary continuity in the microbiomes of these two groups.
5. **Abundance of fiber-digesting microbes:** Saproxylophagous lineages should demonstrate an increased relative abundance of microbes specialized in lignocellulose digestion.

## Future directions

Saproxylophagy is a textbook example of symbiotic innovation and has been studied for over a century. However, our understanding of how it evolved remains fragmented across different disciplines, which prevents a comprehensive synthesis of insights. However, recent advances in sequencing technology, bioinformatics, and conceptual frameworks like the holobiont theory provide new opportunities for integrating the fields of insect phylogenetics, morphology, enzymology, physiology, and microbial ecology. These developments, along with comparative studies of insects offer a promising avenue to investigate the origins of saproxylophagy with a focus on the complex interplay of morphological, physiological, microbial, and ecological factors.

Our observations suggest that future investigations of the origins of saproxylophagy must consider the distribution of saproxylophagous characters in the detritivorous relatives of wood-feeding clades. Taking advantage of the multiple origins of saproxylophagy in insects could facilitate the identification of convergent evolutionary adaptations, i.e. key innovations that led to the evolution of saproxylophagy. Among the many trophic innovations accompanying saproxylophagy, the gut microbiome is the most plastic and metabolically flexible within the lifespan of an insect. This exceptional phenotypic plasticity fosters the enrichment of fiber-digesting microbial symbionts in response to dietary changes. Consequently, the gut microbiome likely confers transient advantages upon protosaproxylophagous lineages while exerting profound and enduring influences on the evolutionary trajectory of saproxylophagy.

## Data availability

The gut microbiome data generated as part of this study are publicly accessible in the NCBI Sequence Read Archive (SRA) under the project number PRJNA1034670.

## Acknowledgments

We extend our sincere gratitude to Dr. Robert R. Dunn for his thoughtful comments on our manuscript. Funding was provided, in part, by NSF Dimensions of Biodiversity project DEB-2030345 to BMW, AM, and CBB

