## Supplementary information for "Chewing Through Challenges: Exploring the Evolutionary Pathways to Wood-Feeding in Insects"

**Supplementary Methods**

**Phylogenetic analysis of Christensenellaceae R-7**

For the phylogenetic analysis of Christensenellaceae R-7 (Fig. 2), sequences representing different levels of the phylogenetic hierarchy were carefully selected, with a focus on standard strains from the American Type Culture Collection (ATCC) for both ingroup and outgroups. After excluding potentially chimeric sequences with pintail scores below 50 and reducing redundancy through OTU selection at 97% identity using Vsearch, alignments from this study and the Silva database were analyzed with IQTREE2, including model selection (-m MFP) and 1,000 Ultrafast Bootstrap replicates for branch support. For more details, please see Schwarz et al. 2023. The alignment comprised 1752 sequences with 1589 columns, including 1459 parsimony-informative sites, 112 singleton sites, and 18 constant sites.

**Microbiome analysis**

16S rRNA gene datasets were downloaded from representative insects (please see Table S1 for accession numbers). Along with the libraries generated from *Micromalthus debilis* and *Epiphragma solatrix*, these published sequence datasets were processed and analyzed as described before (see Schwarz et al., 2023).

To generate the Sankey diagram, Bray-Curtis distances were first calculated using the 'vegdist' function in R, quantifying the similarities in gut microbiome structure across three primary insect diet categories: litter and humus feeders (Column 1 in Fig. 3), wood feeders (Column 2), and non-lignocellulose feeders (Column 3). These calculated distances were then imported into the online SankeyMATIC tool (<https://sankeymatic.com/>) for visualization.

**Supplementary Results**

**Table S1. Relative Abundance of Bacterial Taxa.** This interactive Excel spreadsheet displays the relative abundance of various bacterial taxa identified in 16S rRNA amplicon libraries. It includes data generated from this study alongside data from previously published studies.

**File S1. 16S rRNA Alignment for Christensenellaceae R7 Phylogenetic Analysis.** This fasta file is an alignment of the Christensenellaceae R7 clade, spanning 1590 positions (including gaps), which was used to construct the phylogenetic tree (File S2; Fig. 2).

**File S2. Newick Format Phylogenetic Tree of Christensenellaceae R7.** This Newick file presents the phylogenetic tree derived from the 16S rRNA sequence alignment (File S1) of the Christensenellaceae R7 clade using IQ-TREE.
